# The RED scheme: Rate-constant estimation from pre-steady state weighted ensemble simulations

**DOI:** 10.1101/453647

**Authors:** Alex J. DeGrave, Anthony T. Bogetti, Lillian T. Chong

## Abstract

We present the Rate from Event Durations (RED) scheme, a new scheme that more efficiently calculates rate constants using the weighted ensemble path sampling strategy. This scheme enables rate-constant estimation from shorter trajectories by incorporating the probability distribution of event durations, or barrier crossing times, from a simulation. We have applied the RED scheme to weighted ensemble simulations of a variety of rare-event processes that range in complexity: residue-level simulations of protein conformational switching, atomistic simulations of Na^+^/Cl^−^ association in explicit solvent, and atomistic simulations of protein-protein association in explicit solvent. Rate constants were estimated with up to 50% greater efficiency than the original weighted ensemble scheme. Importantly, our method accounts for systematic error when using data from the entire simulation. The RED scheme is relevant to any simulation strategy that involves unbiased trajectories of similar length to the most probable event duration, including weighted ensemble, milestoning, and standard simulations as well as the construction of Markov state models.

## I. Introduction

Of great interest to chemical physics and biophysics is the estimation of rate constants for long-timescale processes. These rate constants may be directly obtained from molecular simulations with enhanced sampling approaches that maintain rigorous kinetics. Among these approaches are path sampling strategies, which focus the computing power on the functional transitions between stable states rather than the stable states themselves,^1^ exploiting the fact that for rare events, the event duration tb, or barrier crossing time, is much shorter than the associated waiting times between events (tb ≪ k^−1^ where k is the corresponding rate constant).^2,3^ Path sampling strategies fall broadly into two categories: (i) methods that generate continuous transition paths *(e.g.* weighted ensemble^4,5^ and other “splitting” strategies,^6-8^ transition interface sampling,^9^ and forward flux sampling^10,11^), and (ii) methods that generate discontinuous paths (*e.g.* milestoning^12^ and weighted ensemble milestoning^13^). Alternatively, Markov State Models^14,15^—discrete state kinetic models—can be constructed at the post-simulation stage to obtain long-timescale information from either continuous trajectories (*e.g.*, from weighted ensemble simulations)^16,17^ or short, discontinuous trajectories (*e.g.* from adaptive sampling^7^).

One challenge of the weighted ensemble (WE) strategy has been the estimation of rate constants from trajectory ensembles that have not yet reached a steady state. To tackle this challenge, history-augmented Markov State Models that employ “micro-bins” have been applied to estimate rate constants from pre-steady state trajectories.^16,17^ Alternatively, the non-Poisson kinetics of the transient “ramp-up time”—or approach to steady state—of a WE simulation can be incorporated into the rate-constant estimation, improving on previous WE studies of complex biological processes such as large-scale protein conformational transitions^18^ and proteinligand binding^19-21^ that have focused on only the latter portions of the simulations where the rate-constant estimate was no longer sensitive to the earliest (and least probable) successful pathways.

Here we present the Rate from Event Durations (RED) scheme: a more efficient scheme for estimating rate constants that exploits the ramp-up time from the early part of a WE simulation by incorporating the distribution of event durations (barrier-crossing times) that have been sampled. To illustrate the rationale of the RED scheme, we make an analogy of rare-event sampling to a cross-country race in which officials wish to estimate the average rate for runners to surmount the first hill, or barrier (**Figure 1A**). Rather than waiting for all of the runners to complete the race, the officials can estimate the average rate more quickly by constructing a probability distribution of event durations that is solely based on the initial pack of runners that make it over the barrier. The effectiveness of this scheme therefore depends on the extent to which the initial distribution of event durations reflects the width and steepness of the barrier after all runners have finished the race.

**Figure 1.**
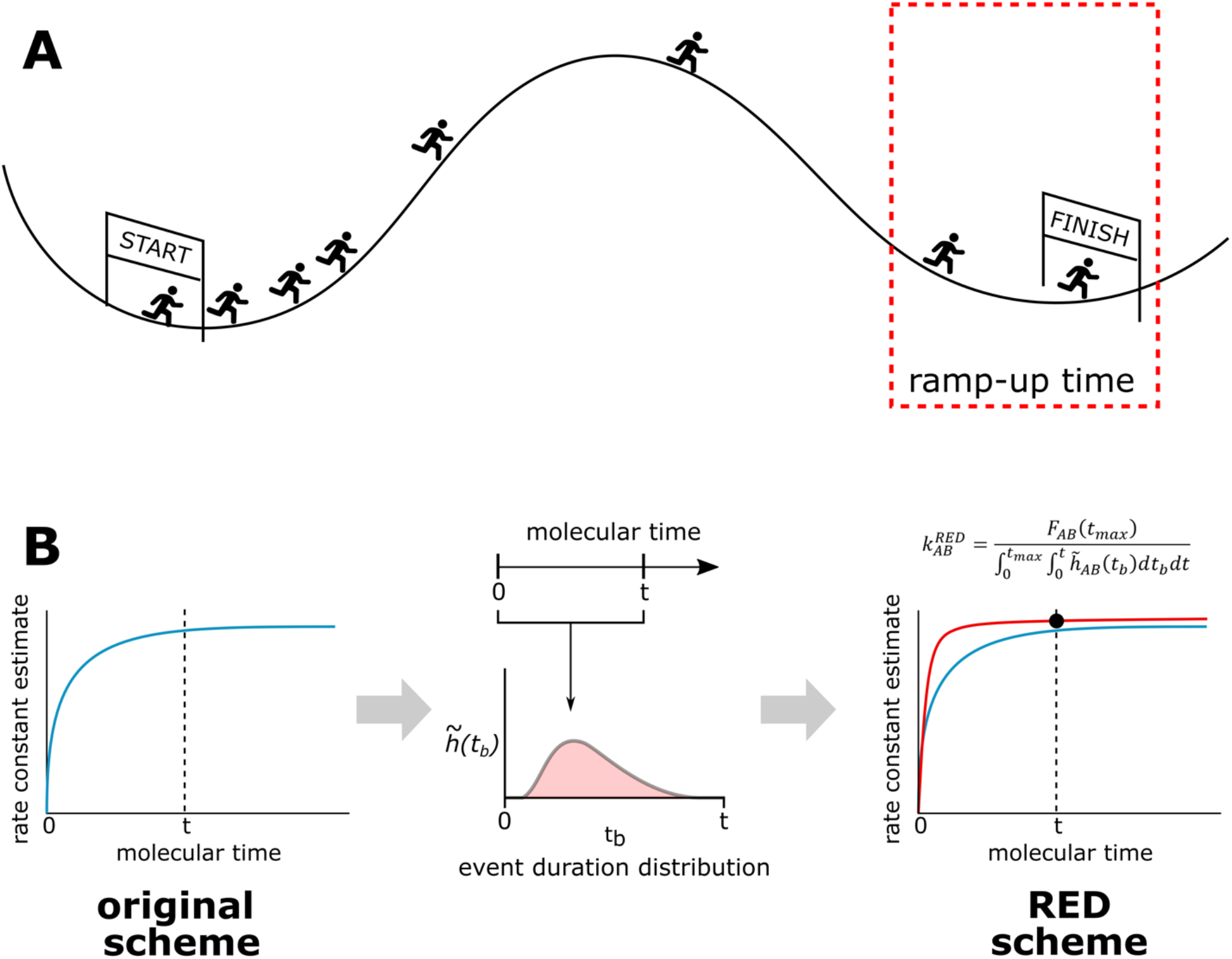
Illustration of the RED Scheme for rate-constant estimation. A) Estimating the rate constant from the ramp-up time is analogous to estimating the average rate at which all runners in a race reach the finish line from the first few finish times. B) In the context of a WE simulation, the RED scheme enhances the efficiency of rate-constant estimation by using the “ramp-up time” for the rate constant, *i.e.* an initial portion of the distribution of event durations. To compare the RED scheme against previous calculation methods, we ask “for a given time *t*, what is the best estimate that our scheme could have produced if we stopped the simulations at time *t*?”

The RED scheme is relevant to any simulation strategy that relies on unbiased pathways of similar length to the typical event duration, including weighted ensemble,^4,5^ milestoning,^12^ and standard simulations as well as the construction of Markov state models.^14,15^ To demonstrate the power of the RED scheme for calculating rate constants, we applied the strategy to a set of three increasingly complex rare-event processes.

First, we applied the RED scheme to residue-level simulations of a protein conformational switching process of an engineered protein-based Ca^2+^ sensor. These simulations have enabled the rational enhancement of the sensor’s response time by as much as 32-fold.^18^ This sensor was engineered using the alternate frame folding (AFF) scheme, fusing together the wild-type calbindin protein and a circular permutant of calbindin such that the two proteins partially overlap in sequence in the resulting calbindin-AFF construct and therefore fold in a mutually exclusive manner.^22^ Importantly, WE simulations of this switching process are an ideal “proof-of-principle” application of the RED scheme as the simulations each exhibit a large “ramp-up time” before steady-state convergence of the rate constant and each simulation captures the entire distribution of event durations.^18^

Second, we applied the RED scheme to the molecular association of Na^+^ and Cl^−^ ions in explicit solvent. This association process was one of four benchmark applications in a previous study that demonstrated the efficiency of WE relative to standard simulations in generating rate constants and pathways.^23^

Finally, we applied the RED scheme to atomistic simulations of a complex biological process in explicit solvent: protein-protein binding. In particular, we re-analyzed a previously completed protein-protein binding simulation that has yielded rate constants and pathways for the barnase and barstar proteins^20^ using <1% of the total simulation time used for a Markov State Model study of the same binding process.^24^

## II. Theory

For a rare-event process, the majority of event durations (barrier crossing times) will be short compared to the waiting times between events. As the system evolves in time and begins to generate event duration times that are substantially longer than the most probable event duration, the distribution of waiting times becomes near-exponential, which is consistent with a Poisson point process in which the events are stochastic and independent.^25^ However, when the simulations of a rare-event process are only as long as the most probable event duration—as is often the case for WE and other rare-events sampling strategies—the number of events per unit time displays transient, pre-steady state behavior, and the initial edge of the distribution of waiting times deviates from an exponential distribution. Our Rates from Event Durations (RED) scheme leverages this transient behavior to estimate rate constants from pre-steady-state trajectories. Below, we briefly summarize the weighted ensemble (WE) strategy and then present details of the original WE scheme for rate-constant estimation and the RED scheme.

### A. The weighted ensemble (WE) strategy

The WE strategy enhances the sampling of rare events by orchestrating the periodic resampling of parallel, weighted trajectories.^4^ The goal of the strategy is to provide reasonably even coverage of configurational space – typically divided into bins along a progress coordinate toward the target state – to yield an ensemble of continuous, successful pathways with rigorous kinetics. The resampling step is performed at a fixed time interval *τ* and involves evaluating trajectories in the same bin for either replication or combination to maintain the same number of target trajectories/bin. Rigorous management of trajectory weights ensures that no bias is introduced into the dynamics. To maintain non-equilibrium steady-state conditions, trajectories that reach the target state are “recycled”, *i.e.* terminated followed by initiation of a new trajectory with the same weight.

### B. Original WE scheme for rate-constant estimation

In the original WE scheme, the macroscopic rate constant *k_AB_* for a rare-event process involving an initial state A and target state B is computed as follows:^26^

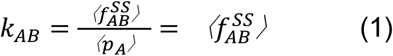

where 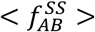 is the running average of the conditional flux of probability carried by trajectories originating in state A and arriving in state B and <*p_A_>* is the running average of the fraction of trajectories more recently in A than in B, which is equal to one in non-equilibrium steady-state WE simulations. In practice, if a steady state has not been reached, then 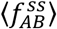 is approximated by the running average ⟨*f_AB_*⟩ of the conditional flux (not necessarily steady state) from state A to state B. For bimolecular processes, we divide equation (1) by the effective molar concentration C_0_ of the associating molecules to estimate a rate constant in units of M^−1^s^−1^.

### C. Rate from Event Durations (RED) scheme

The Rate from Event Durations (RED) scheme reduces the impact of transient effects from a WE simulation on rate-constant estimation by incorporating the distribution of sampled event durations (barrier crossing times *t_b_* which exclude the dwell time in state A). The motivation behind this scheme is that short WE simulations may not capture pathways with relatively long barrier-crossing times that have yet to enter state B; therefore, the original WE scheme tends to underestimate the true rate constant by a predictable quantity that depends on the probability of observing pathways with longer event durations. The RED scheme incorporates this quantity as a correction factor to the rate-constant estimate of the original scheme at a given time of the simulation.

We consider a rare-event process with the following properties:

1. The system is in an initial state A at time *t =* 0 such that an event of duration *t_b_* is less than or equal to the longest possible trajectory length *t_max_* of the WE simulation.
2. While in the initial state A, the system has a constant probability per unit time of initiating successful transition path to the target state B, denoted *k_AB_*.
3. The event durations are assumed to be randomly distributed according to a probability density function *h_AB_*, where 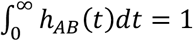.
4. Upon arriving in a target state B, the system is immediately “recycled” to the initial state A.

To derive an expression for estimating the rate constant, we begin by defining the flux *f_AB_* from an initial state A into a target state B as a convolution of the rate constant *k_AB_* for completing the A→B transition in a time *t_b_* distributed according to *h_AB_* (see Supporting Information for additional details):

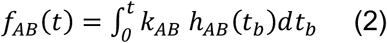

We then integrate and rearrange equation (2) to obtain an expression for *k_AB_* that depends only on the true cumulative number of events *f_AB_ (t_max_)* and cumulative distribution of event durations *H_AB_(t)*:

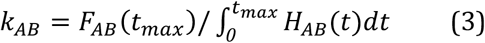

where the numerator 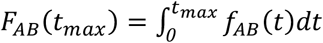 and the denominator is the integral of *H_AB_ (t)* over all values of *t* ranging from 0 to *t_max_* where 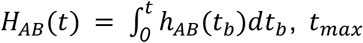, and *h_AB_(t_b_)* is the true distribution of event durations. Compared with the original WE scheme, where the denominator would be the time *t_max_*, the denominator in equation (3) represents a “corrected time”, which accounts for the time during which it was *possible* to see events. Equivalently, the denominator in equation (1) of the original WE scheme could be written as 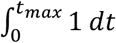, which indicates that an estimate derived from equation (3) would be greater than that of the original WE scheme, since *H_AB_(t)* is a cumulative density function that is less than one.

Next, we use equation (3) to derive an estimate for the rate constant based on the “observed” distribution of event durations that are sampled by the WE simulation. While we may naively estimate *h_AB_(t_b_)* as the observed histogram 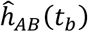 of event durations, the observed histogram 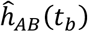 is likely skewed toward shorter event durations due to the transient phase for the time evolution of the rate constant 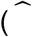 indicates the observed quantity). To obtain a corrected estimate 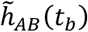 of the histogram, we divide the observed histogram 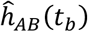 by the interval of time *(t_max_ - t_b_)* in which it is possible to observe an event of duration *t_b_* from a simulation with a maximum trajectory length *t_max_*:

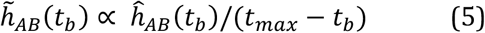

where the constant of proportionality is chosen such that the corrected 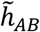 is normalized 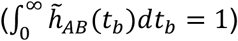.

Finally, we define the RED scheme estimate 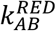 of the true rate constant *k_AB_* as follows:

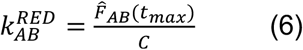

where 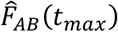 is the observed cumulative probability of *A → B* transitions up to the maximum trajectory length *t_max_;* and the denominator is a correction factor *C* equal to 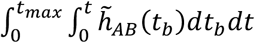 in units of time, yielding a rateconstant estimate 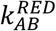 in units of inverse time. For bimolecular processes, we divide equation (6) by the effective molar concentration *C_0_* of the associating molecules to estimate a rate constant in units of M^−1^s^−1^ (as is also the case for the original WE scheme).

### D. Error estimation for rate constants

In cases where it is not possible to sample the entire distribution of event durations, the RED scheme provides a framework for understanding the error that results from not observing trajectories with longer event durations. Given a maximum trajectory length *t_max_*, the corrected estimate 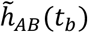 of the event duration distribution *h_AB_(t_b_*) will be zero for *t_b_> t_max_* and, since 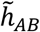 is normalized such that 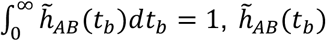 will be artificially inflated for *tb < t_max_*:

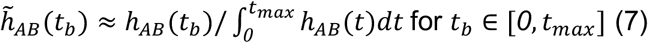

If we plug the right-hand side of equation (7) back into equation (6), we find that 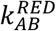 underestimates *k_AB_* by a factor of 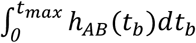, the observed fraction of the distribution of event durations:

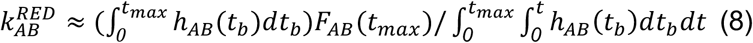

For example, if 20% of pathways reaching the target state have longer event durations *t_b_* than the maximum trajectory length *t_max_* and are therefore not captured during the simulation, then we tend to underestimate the true rate constant *k_AB_* by 20%. Despite this underestimation, the RED scheme estimate is still an improvement over the original scheme for estimating rate constants (equation (1)).

For multiple, independent WE simulations *1,2,…,N*, we estimated uncertainties in the rate constants by first applying the RED scheme individually to map each simulation *i.* to a corresponding rate constant estimate *k^RED,i^*, and then applying Bayesian bootstrapping^27^ to estimate 95% credibility regions (CR). To prevent underestimating the uncertainty, the distributions of event durations 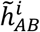 were calculated independently for each simulation, as pooling data to make a smoother estimate of *h_AB_* would introduce correlations and therefore break the independence between the *k^RED,i^*. For cases where only a single WE simulation was run *(i.e.* for barnase-barstar association), the uncertainty in the rate constant calculated by the RED scheme is not reported as the error estimation is not straightforward in these cases (see Supporting Information).

## III. Methods

### A. WE simulations

All WE simulations were run using the open-source, highly scalable WESTPA software package (https://westpa.github.io/westpa).^28^ WE parameters and details of dynamics propagation are provided below for each rare-event process.

#### Protein conformational switching

As described in DeGrave *et al.^18^* 10 independent WE simulations were previously run to generate N’→N switching pathways of the wild-type E65’Q calbindin-AFF construct under nonequilibrium steady-state conditions. Each WE simulation was run for 2000 WE iterations with a fixed time interval *τ* of 100 ps and a target number of 5 trajectories/bin, yielding an aggregate simulation time of 65 μs for each simulation. A two-dimensional progress coordinate was defined as (i) the pseudo-atom RMSD of the N frame after aligning on the folded N frame structure, and (ii) the pseudo-atom RMSD of the N’ frame after aligning on the folded N’ frame. Dynamics were propagated using a Brownian dynamics algorithm with hydrodynamic interactions, as implemented in the UIOWA-BD software.^29,30^ All analysis was performed with conformations sampled every 50 ps. A minimal residue-level protein model was employed in which each residue is represented by a single pseudo-atom at the position of its *C_α_* atom. The conformational dynamics of the protein were governed by a Gō-type potential energy function^31,32^ that was parameterized to reproduce the experimental folding free energies of the isolated wild-type protein and circular permutant of the protein.^18^

#### Na^+^/Cl^−^ association

Five independent WE simulations were run to generate pathways of the Na^+^/Cl^−^ association process under non-equilibrium steady-state conditions. Each WE simulation was run for 1000 WE iterations with a fixed time interval *τ* of 2 ps for each iteration and a target number of 4 trajectories/bin, yielding an aggregate simulation time of 0.2 μs for each simulation. A one-dimensional progress coordinate was defined as the distance between the Na^+^ and Cl^−^ ions; bins were placed every 1 Åfrom a separation distance of 12 Å (unassociated state) to 2.6 Å (associated state). Dynamics were propagated using the AMBER18 software package^33^ with the TIP3P water model^34^ and corresponding Joung and Cheatham parameters for the Na^+^ and Cl^−^ ions.^35^ Simulations were started from an unassociated state with a 12-Å separation between the Na^+^ and Cl^−^ ions and a sufficiently large truncated octahedral box of explicit water molecules to provide a minimum 12 Å clearance between the ions and box walls, yielding an effective ion concentration C_0_ of 2.8 mM. Temperature and pressure were maintained at 298 K and 1 atm using the Langevin thermostat (collision frequency of 1 ps^−1^) and Monte Carlo barostat (with 100 fs between attempts to adjust the system volume), respectively. Non-bonded interactions were truncated at 10 Å and long-range electrostatics were treated using the particle mesh Ewald method.^36^

#### Protein-protein association

As described in Saglam and Chong,^20^ a single WE simulation was previously run to generate pathways of the association process of the barnase and barstar proteins under equilibrium conditions.^20^The WE simulation was run for 650 WE iterations with a fixed time interval *τ* of 20 ps for each iteration and a fixed total number of 1600 trajectories at all times during the simulation, yielding an aggregate simulation time of 18 μs. A two-dimensional progress coordinate was defined as (i) the minimum separation distance between barnase and barstar, and (ii) a “binding” RMSD, which was determined by first aligning on barnase in the crystal structure of the barnase-barstar complex^37^ and then calculating the heavy-atom RMSD of barstar residues D35 and D39. Dynamics were propagated using the Gromacs 4.6.7 software package^38^ with the Amber ff03* force field^39^, TIP3P water model^40^, and corresponding Joung and Cheatham ion parameters.^35^ The system was immersed in a sufficiently large dodecahedron box of explicit water molecules to provide a minimum 12 Å clearance between the solutes and box walls for the unbound states in which the binding partners were separated by 20 Å. A total of 31 Na^+^ and 29 Cl^−^ ions were included to neutralize the net charge of the protein system and to yield the experimental ionic strength (50 mM).^41^ The entire simulation system consisted of ~100,000 atoms with an effective protein concentration C_0_ of 1.7 mM. Heavy-atom coordinates for initial models of the unbound proteins were extracted from the crystal structure of the barnase-barstar complex (PDB code: 1BRS).^37^

### B. Standard simulations

To validate the rate constants computed from the WE simulations for the protein conformational switching process and Na^+^/Cl^−^ association process, an extensive set of standard simulations was run to provide “gold standard” rate constants for comparison. Given the computationally prohibitive timescales for the barnase-barstar association process, no standard simulations were run for this process; instead, the experimental association rate constant is used to validate the computed association rate constant from the WE simulation. For the protein conformational switching process, 50 2-μs standard simulations were run. For the Na^+^/Cl^−^ association process, 10 1-μs standard simulations were run. Dynamics were propagated as described above for the corresponding WE simulations.

## IV. Results and Discussion

We have developed the Rate from Event Durations (RED) scheme: a new scheme for rate-constant estimation that reduces the impact of transient effects by using the distribution of event durations that correspond to simulated pathways of the rare event. To demonstrate the effectiveness of the RED scheme, we have applied the scheme to simulations of three rare-event processes: (i) residue-level simulations of protein conformational switching by an engineered protein-based calcium sensor; (ii) atomistic simulations of Na^+^/Cl^−^ association in explicit solvent; and (iii) atomistic simulations of protein-protein association in explicit solvent. The effectiveness of the RED scheme was evaluated by monitoring the time-evolution of the rate constant, incorporating the distribution of event durations up to each time point (**Figure 1B**).

### A. Application to residue-level simulations of protein switching

The switching process of the engineered calbindin-AFF system (**Figure 2A**), as simulated using a residue-level model, is an example of a case where the RED scheme would be expected to be particularly effective in enabling the calculation of rate constants from shorter trajectories. This expectation is based on the relatively long “ramp up time” of the flux into steady state from a given WE simulation.

**Figure 2.**
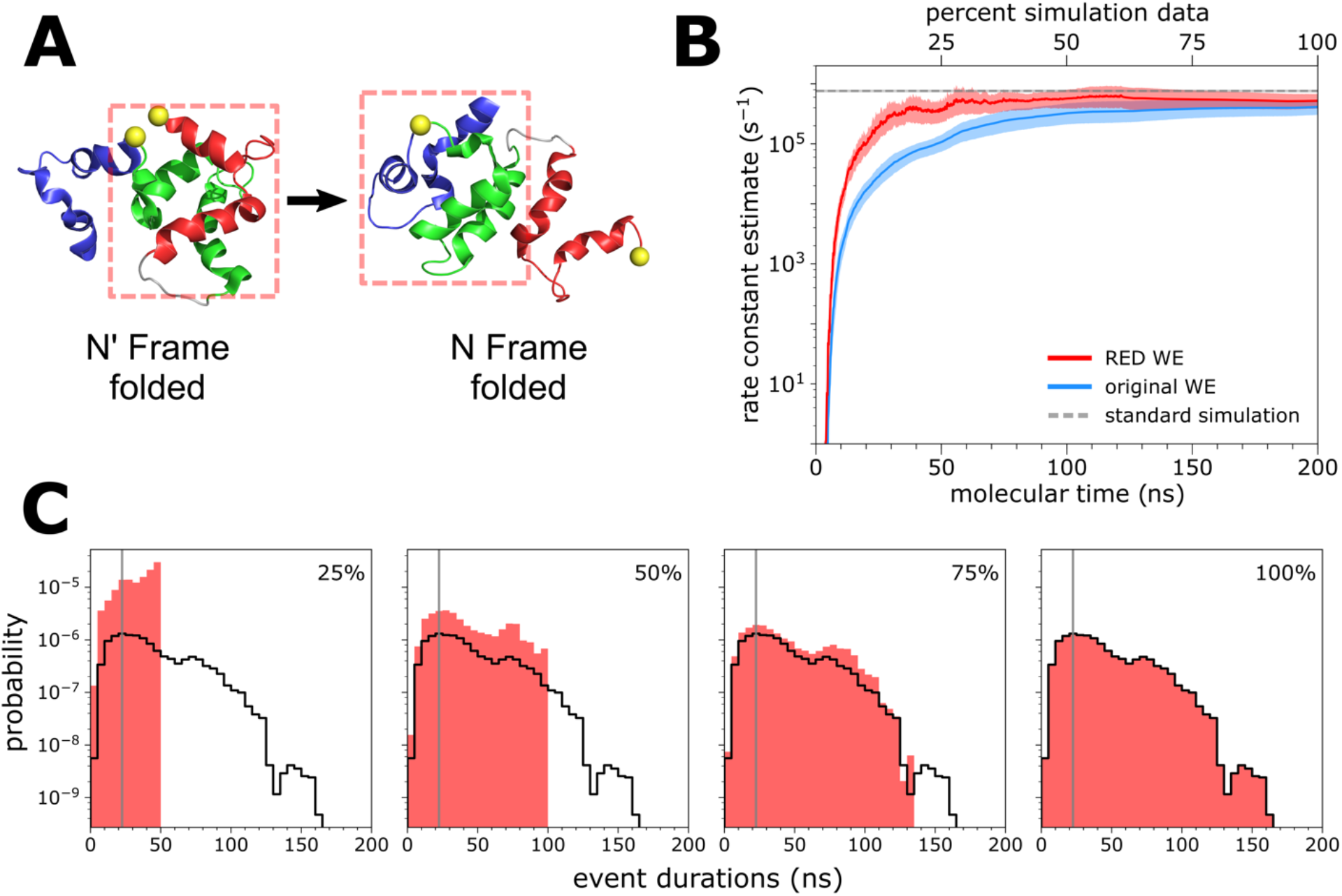
Residue-level simulations of protein conformational switching. A) A schematic of the calbindin-AFF switch, showing the initial N’ state and target N state of the simulated switching process. B) Comparison of the N’→N switching rate constant (average of 10 WE simulations) using the original WE scheme 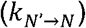 and RED scheme 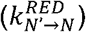, as a function of molecular time, or Nτ where N is the number of WE iterations and *τ* is the fixed time interval (100 ps in this case) of each WE iteration. See also **Table S1**. Ten WE simulations were run for each scheme. The RED scheme was applied using the first 25%, 50%, and 75% from each WE simulation. Also shown is the rate constant calculated from 50 2-μs standard simulations (horizontal dashed line). The shaded regions show the nominal 95% credibility regions (CR) as a function of molecular time from Bayesian bootstrapping;^27^ the CR from standard simulations is displayed, but too small to be visible. C) Estimates of the probability density function hAB of event durations for the switching process, as sampled by the first 25%, 50%, and 75% of a representative WE simulation. The vertical gray line indicates the most probable event duration based on the distribution from 100% of the simulation (delineated in black).

To determine the effectiveness of the RED scheme, we examined the evolution of the rate constant 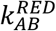 as a function of the molecular time, where at any given time the estimate 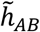 is based only on data from all 10 independent WE simulations that were generated up to and including that time. The RED scheme yields faster convergence of the rate constant 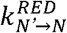 for the N’→N switching process (**Figure 2B**), requiring only the first 25% of the WE simulation data to reproduce the rate constant from standard simulations (50 2-μs simulations). This is almost 50% more efficient than the original scheme, which only began to converge after 75% of the simulation data had been collected and underestimated the rate constant by a factor of two (compared with that from standard simulations) due to the slow transient phase.

We determined the extent of simulation required for estimating rate constants by monitoring the position of the maximum in the distribution of event durations. If the position did not shift substantially--meaning that the most probable event duration reached a consistent value--we considered the simulation as being converged for the purpose of estimating rate constants using the RED scheme. **Figures 2B** and **2C**show that the most probable event duration (as defined from 100% of the data collected) is captured within the initial 25% of a given WE simulation; furthermore, the cumulative probability distribution of event durations is well-resolved and not skewed towards short values, with low probability events occurring consistently throughout the course of the simulation.

### B. Application to atomic-level simulations of Na^+^/Cl^−^ association

Na^+^/Cl^−^ association in explicit solvent (**Figure 3A**) occurs on the ns timescale, which is orders of magnitude faster than the calbindin-AFF switching process and the complex processes that follow. Given the fast event durations of the ion-pair association, it is not expected that the RED scheme would provide much benefit over standard WE rate constant estimation. We found that this was indeed the case, as the system does not exhibit a “rampup time” (**Figure 3B**) and the most probable event duration is sufficiently sampled with less than 25% of the data collected (**Figure 3C**).

**Figure 3.**
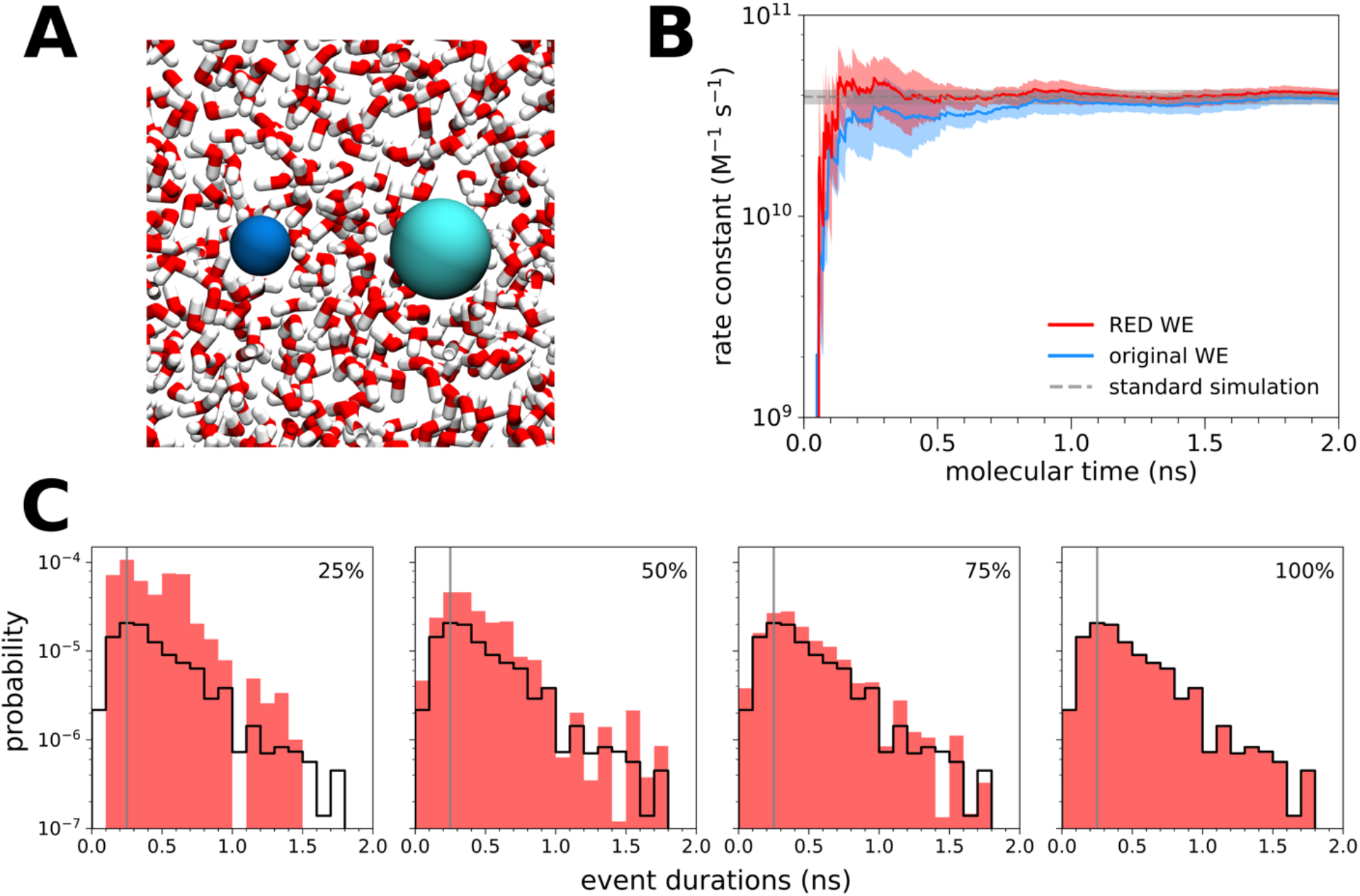
Atomistic simulations of Na^+/^Cl^−^ association in explicit solvent. A) The Na^+^/Cl^−^ system in explicit solvent. **B)**Comparison of the Na^+^/Cl^−^ association rate constant 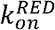 (average of five WE simulations) using the original WE and RED schemes, plotted as a function of molecular time, or Nτ where N is the number of WE iterations and *τ* is the fixed time interval (2 ps) of each iteration. See also **Table S1**. Five WE simulations were analyzed with each scheme. The RED scheme was applied using the first 25%, 50%, and 75% from each WE simulation. Also shown is the rate constant calculated from 10 1-μs standard simulations (horizontal dashed line). The shaded regions show the nominal 95% credibility regions (CR) as a function of molecular time from Bayesian bootstrapping.^27^ The y-axis shown is only a portion of the full set of values in order to more clearly show the comparison between the two schemes; see **Figure S1** for a plot using the full range of data. C) Estimates of the probability density function hAB of event durations for the molecular association process, as sampled by the first 25%, 50%, and 75% of the 5 WE simulations. The vertical gray line indicates the most probable event duration based on the distribution from 100% of the simulation (delineated in black).

### C. Application to atomistic simulations of long-timescale processes in explicit solvent

To test the effectiveness of the RED scheme in estimating rate constants from more detailed simulations of complex biological processes, we applied the scheme to a single WE simulation of a protein-protein binding process. This simulation involved the diffusion-controlled association of the barnase and barstar proteins using atomistic protein models with explicit solvent (**Figure 4A**). While this simulation was not performed with recycling enabled and therefore violates one of the RED scheme’s assumptions, based on the extremely short length of the simulation compared to the mean first passage time, the weight of the trajectories that would have been recycled is extremely low, such that negligible inaccuracy is introduced. When applied to this simulation, the RED scheme is at least 25% more effective than the original scheme in estimating rate constants given that the WE simulation has just finished ramping up to a steady state. Similar to the simulation of protein conformational switching, this simulation exhibits a long “ramp-up-time” (**Figure 4B**). In contrast, the most probable event duration is relatively long (6 ns) and just shy of being captured within the first 50% of the simulation, underestimating the rate constant compared to the eventual converged value (**Figure 4C**). Based on the first 75% of the simulation, the rate-constant is still underestimated, but due to another reason: the most probable event duration is actually *longer* than that based on the entire simulation. Due to the large size of the simulation system (~100,000 atoms) and the relatively long timescales of the protein-protein binding process, only one WE simulation was carried out; therefore, no error analysis was performed for the estimated rate constants.

**Figure 4.**
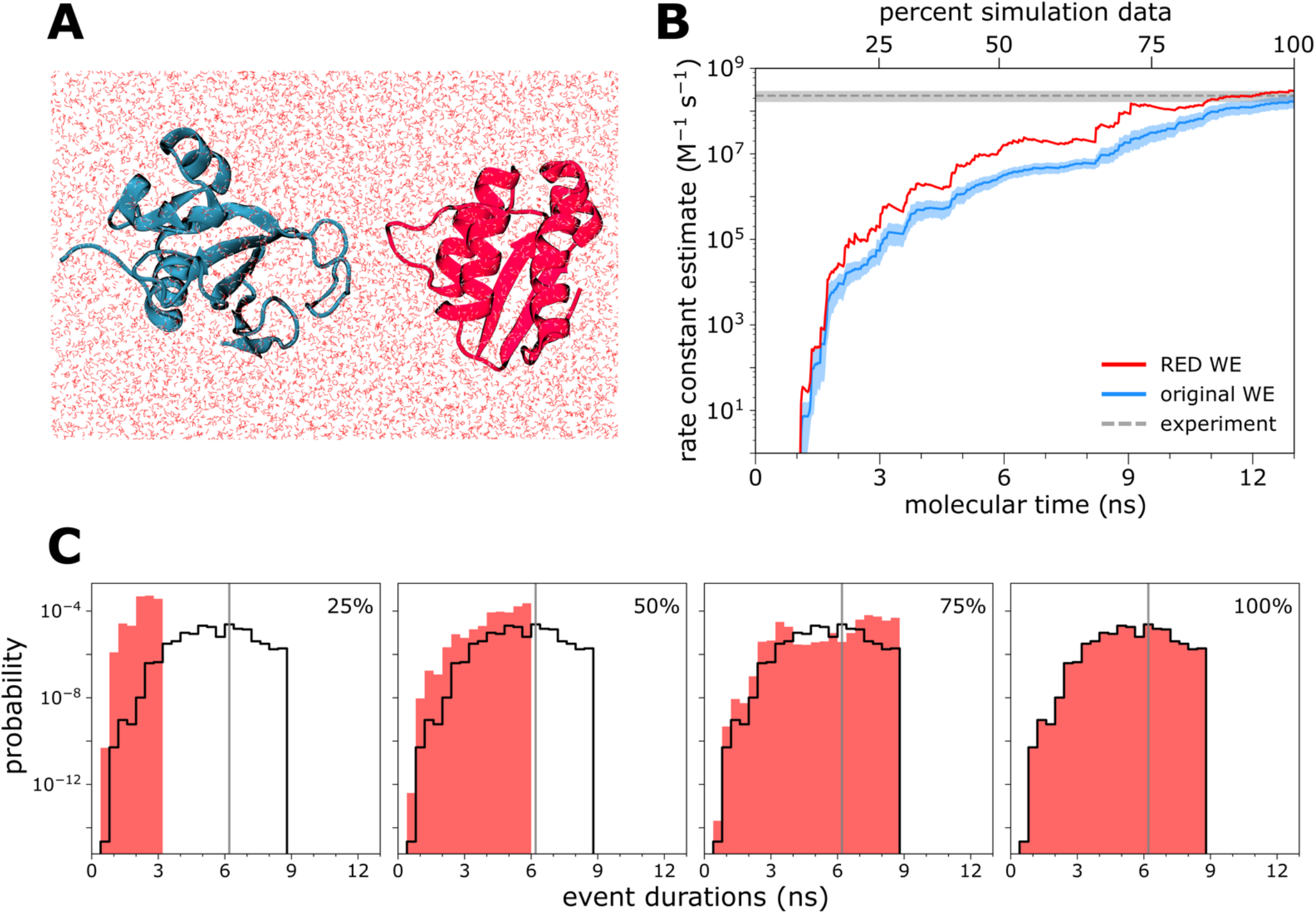
Atomistic simulations of protein-protein association in explicit solvent. A) A representative unbound state of the barnase and barstar proteins in explicit solvent. B) Comparison of the barnase-barstar association rate constant 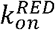 using the original WE and RED schemes, plotted as a function of molecular time, or Nτwhere N is the number of WE iterations and *τ* is the fixed time interval (20 ps) of each WE iteration. See also **Table S1**. A single WE simulation was analyzed with each scheme. To test the length of simulation required for a converged rate-constant estimation, the RED scheme was applied using the first 25%, 50%, and 75% from each WE simulation. Also shown is the rate constant from experiment (horizontal dashed line); the uncertainty (shaded gray) is the 95% confidence interval determined from standard errors of the mean reported for the experimental results;^41^ the uncertainty of the rate constant from the original WE scheme is the 95% confidence interval by Monte Carlo bootstrapping.^42^ C) Estimates of the probability density function hAB of event durations for the protein-protein association process, as sampled by the first 25%, 50%, and 75% of one of the 10 WE simulations depicted in panel A. The vertical gray line indicates the most probable event duration based on the distribution from 100% of the simulation (delineated in black).

### D. When is the RED scheme effective and how do we monitor convergence?

Regardless of the simulation model resolution, the RED scheme is particularly efficient in rate-constant estimation for rare events that involve long “ramp ups” in the time evolution of the estimated rate constant. For atomically detailed simulations, the RED scheme works well for long-timescale processes on the μs timescale or beyond. In this study, the RED scheme is of great benefit to residue-level simulations of the protein conformational switching process involving the calbindin-AFF switch due to the large ramp-up time in the flux into the target state, and to atomistic simulations of protein-protein binding on the μs timescale. On the other hand, the RED scheme has little impact on the efficiency of rate-constant estimation for the simulations of Na^+^/Cl^−^ association since this process is relatively rapid and does not exhibit a large ramp-up time in the flux into the target, associated state. As recommended for the original WE scheme,^23^ the RED scheme is more likely to yield converged rate constants for a process if the most probable event duration has been sampled. Provided that this is the case, the RED scheme estimates rate constants more efficiently than the original WE scheme.

An effective convergence criterion for determining the amount of simulation data necessary for the RED scheme is to generate a sufficient number of successful events such that the position of the maximum in the distribution of event durations *(i.e.* the most probable value) does not change substantially. For both the calbindin-AFF switching process and Na^+^/Cl^−^ association process, trajectories with the most probable event duration are already sampled within the first 25% of the WE simulation. On the other hand, for the barnase-barstar association process, the most probable event duration begins to stabilize only after 75% of the simulation is completed. If the most probable event duration continues to evolve after completing the simulation, the system is likely far from a steady state and will require generating a much larger number of successful pathways to yield a converged rate-constant estimate. Alternatively, if the event duration distribution involves a long tail, it may be necessary to sample more of the distribution than just the most probable value.

For challenging cases in which a large amount of computing has already been invested, we recommend applying the RED scheme to quickly gauge the extent to which the simulation has reached steady state. If the estimated rate constant is orders of magnitude from that of the expected timescales, then we recommend constructing a history-augmented Markov State Model^43^ to adjust trajectory weights to values more representative of steady state conditions and carrying out a separate WE simulation with the adjusted weights.

## V. Conclusions

We have developed the Rate from Event Durations (RED) scheme: a new scheme for calculating rate constants within the framework of the weighted ensemble (WE) strategy that reduces the impact of transient effects on rate-constant estimation. While the RED scheme does not eliminate the need to observe the substantial portion of the distribution of barrier-crossing times, we anticipate that this scheme—by correctly incorporating the transient phase into the rate-constant estimation rather than “throwing it away”—will enable more accurate estimation of rate constants earlier on in a simulation, using a fraction of the total simulation time required by the original WE scheme. Furthermore, as demonstrated by our results for protein-protein association, the RED scheme could be especially important for estimating the rate constants of challenging biological processes that feature long transient phases. Importantly, the scheme accounts for systematic error when using data from the entire simulation—even before the molecular time exceeds the maximum event duration.

## Dedication

This paper is dedicated to Maud Menten, a Canadian woman who—together with Leonor Michaelis—developed the ground-breaking Michaelis-Menten equation for enzyme kinetics. To work with Michaelis, she crossed the Atlantic by ship in 1912--not long after the *Titanic* sank. Unable to find a faculty position in her native Canada, she joined the faculty in the medical school at the University of Pittsburgh in 1918.

## Data Availability Statement

The data that supports the findings of this study are available within the article and its supplementary material. A Python implementation of the RED scheme for use with the WESTPA software package^8^ is available on GitHub (https://qithub.com/westpa/user_submitted_scripts/tree/main/RED_scheme).

## Acknowledgements

This work was supported by the NIH (1R01GM115805-01) and NSF (CHE-1807301) to L.T.C., the University of Pittsburgh to A.J.D. (Honors College Brackenridge Undergraduate Research Fellowship) and A.T.B. (Arts & Sciences Fellowship). Computational resources were provided by NSF XSEDE allocation TG-MCB100109 to L.T.C., NSF CNS-1229064, and the University of Pittsburgh’s Center for Research Computing. We thank Daniel Zuckerman (OHSU) and Ali Saglam (U. Pittsburgh) for insightful discussions.

The authors declare the following competing financial interest: L.T.C. is an Open Science Fellow with Silicon Therapeutics.

