## Supporting Information for "The RED scheme: Rate-constant estimation from pre-steady state weighted ensemble simulations"

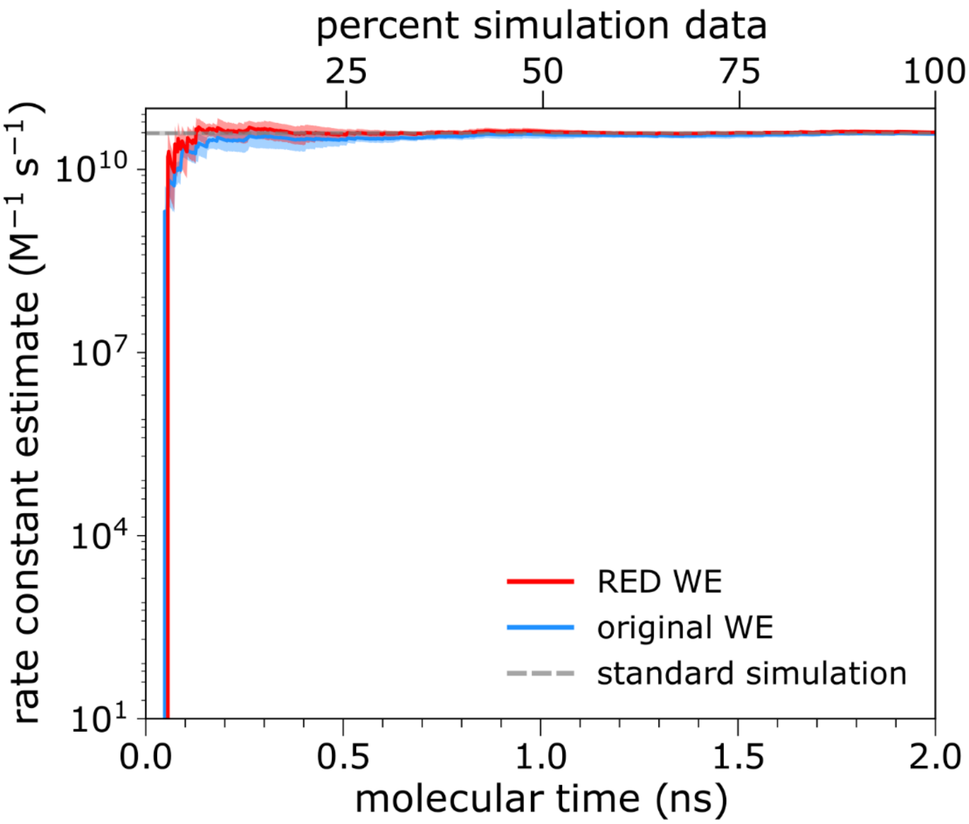


**Figure S1.** Zoomed out view of Figure 3B showing the full range of the y-axis for the time-evolution of the rate-constant estimate from WE simulations of $Na^+/Cl^-$ association.

**Table S1.** Estimated rate constants for the systems studied using the original and RED schemes ($k_{AB}$ and $k_{AB}^{RED}$, respectively). For validation, available estimates from standard "brute force" simulations $k_{AB}$(BF) or experiments $k_{AB}$(EXP) are included in the last column. Uncertainties in the estimated rate constants for calbindin-AFF switching and $Na^{+}$/$Cl^{-}$ association processes are 95% credible regions (CR) by Bayesian bootstrapping;[7] for barnase/barstar binding, uncertainties in estimates from the original scheme are 95% confidence intervals by Monte Carlo bootstrapping[1] and those for the experimental value are also 95% confidence intervals.

| | WE simulations | | Validation |
|---|---|---|---|
| **process simulated** | $\boldsymbol{k_{AB}}$ | $\boldsymbol{k_{AB}^{RED}}$ | $\boldsymbol{k_{AB}}$**(BF/EXP)** |
| **calbindin-AFF switching ($s^{-1}$)** | (4.12 ± 1.10) x $10^{5}$ | (5.22 ± 1.49) x $10^{5}$ | BF: (7.63 ± 0.38) x $10^{5}$ |
| **$Na^{+}$/$Cl^{-}$ association ($M^{-1}s^{-1}$)** | (3.84 ± 0.22) x $10^{10}$ | (4.08 ± 0.20) x $10^{10}$ | BF: (3.93 ± 0.33) x $10^{10}$ |
| **barnase/barstar binding($M^{-1}s^{-1}$)** | (1.67± 0.54) x $10^{8}$ | 2.99 x $10^{8}$ | EXP:[8] (2.30 ± 1.31) x $10^{8}$ |

**Detailed derivation of equation (3)**

To begin, we consider the relationship between the instantaneous flux $f_{AB}(t)$ at time $t$, the rate constant $k_{AB}$, and the true probability distribution $h_{AB}$ of event durations. To be precise, $f_{AB}(t)$ is the time derivative of a cumulative flux function $F_{AB}$, where $F_{AB}(t)$ gives the total number of $A \rightarrow B$ events observed by time $t$.

For an $A \rightarrow B$ transition observed at time $t$ with an event duration of $t_b$, the event must have been initiated at time $t - t_b$. Thus, the instantaneous flux depends on (i) the probability $h_{AB}(t_b)$ that barrier-crossing takes time $t_b$ and (ii) the frequency at which $A \rightarrow B$ events are initiated at time $t - t_b$, which is $k_{AB}$ when $t - t_b > 0$, and zero otherwise, *since the process does not start until time* $0$.

To derive an expression for $f_{AB}(t)$, we integrate over all possible event durations $t_b$. Formally, this is a convolution of $h_{AB}$ with the function that is $k_{AB}$ for parameters greater than zero, and zero otherwise:

$$k(t) \coloneqq \begin{cases} k_{AB}; t \geq 0 \\ 0; t < 0 \end{cases} \qquad \text{(S.1)}$$

$$f_{AB}(t) = k(t) * h(t) = \int_{-\infty}^{\infty} k(t - t_b) h_{AB}(t_b) dt_b \qquad \text{(S.2)}$$

Since both functions in the convolution in equation (S.2) are non-zero:

$$f_{AB}(t) = \int_0^t k_{AB} h_{AB}(t_b) dt_b \qquad \text{(S.3)}$$

Next, we integrate both sides of equation (S.3) with respect to $t$:

$$\int_0^{t_{max}} f_{AB}(t) dt = \int_0^{t_{max}} \int_0^t k_{AB} h_{AB}(t_b) dt_b \, dt \qquad \text{(S.4)}$$

$$F_{AB}(t_{max}) - F_{AB}(0) = k_{AB} \int_0^{t_{max}} \int_0^t h_{AB}(t_b) dt_b \, dt \qquad \text{(S.5)}$$

$$F_{AB}(t_{max}) = k_{AB} \int_0^{t_{max}} H_{AB}(t) dt \qquad \text{(S.6)}$$

We define the cumulative distribution function $H_{AB}$ as the integral of the probability density function $h_{AB}$, that is, $H_{AB}(t) = \int_0^t h_{AB}(t_b) dt_b$. The left-hand side (LHS) of equation (S.5) is given by the definition of $F_{AB}$ and the fundamental theorem of calculus, while the right-hand side (RHS) is given by the fact that $k_{AB}$ does not depend on the parameters $t$ and $t_b$ that are being integrated. The LHS of equation (S.6) results because the number of events $F_{AB}(0)$ observed by $t = 0$ is necessarily zero, while the RHS is given by the definition of $H_{AB}$.

Finally, to obtain equation (3) for $k_{AB}$, we divide both sides by $\int_0^{t_{max}} H_{AB}(t) dt$:

$$k_{AB} = \frac{F_{AB}(t_{max})}{\int_0^{t_{max}} H_{AB}(t) dt}$$

where $F_{AB}(t_{max})$ is the cumulative number of events and the integral $\int_0^{t_{max}} H_{AB}(t) dt$ is in units of time, yielding a rate constant $k_{AB}$ that has units of inverse time.

### Challenges of estimating uncertainties from a single WE simulation

For a single WE simulation, we are unaware of a method for estimating uncertainties in rate constants obtained using the RED scheme. While previous studies have used Monte Carlo block bootstrap approaches[1] to estimate uncertainties in rate constants obtained using the original scheme,[2,3] such block bootstrap approaches generally pertain to stationary processes[4] and have therefore only focused on the latter portions of the simulation when the flux $f_{AB}$ could be assumed to be stationary. Although bootstrap approaches have been formulated for certain non-stationary processes,[5,6] none of these approaches appear to be applicable to RED scheme, which exploits the transient, non-stationary phase of $f_{AB}$ for more efficient rate-constant estimation. Exploration of bootstrap approaches that perform reliably during the transient phase of simulations remains an area for future work.
